# Identifying task-relevant spectral signatures of perceptual categorization in the human cortex

**DOI:** 10.1101/483487

**Authors:** Ilya Kuzovkin, Juan R. Vidal, Marcela Perrone-Bertlotti, Philippe Kahane, Sylvain Rheims, Jaan Aru, Jean-Philippe Lachaux, Raul Vicente

## Abstract

Human brain has developed mechanisms to efficiently decode sensory information according to perceptual categories of high prevalence in the environment, such as faces, symbols, objects. Neural activity produced within localized brain networks has been associated with the process that integrates both sensory bottom-up and cognitive top-down information processing. Yet, how specifically the different types and components of neural responses reflect the local networks selectivity for categorical information processing is still unknown. By mimicking the decoding of the sensory information with machine learning we can obtain accurate artificial decoding models. Having the artificial system functionally on par with the biological one we can analyze the mechanics of the artificial system to gain insights into the inner workings of its biological counterpart. In this work we train a Random Forest classification model to decode eight perceptual categories from visual stimuli given a broad spectrum of human intracranial signals (4 – 150 Hz) obtained during a visual perception task, and analyze which of the spectral features the algorithm deemed relevant to the perceptual decoding. We show that network selectivity for a single or multiple categories in sensory and non-sensory cortices is related to specific patterns of power increases and decreases in both low (4 – 50 Hz) and high (50 – 150 Hz) frequency bands. We demonstrate that the locations and patterns of activity that are identified by the algorithm not only coincide with the known spectro-spatial signatures, but extend our knowledge by uncovering additional spectral signatures describing neural mechanisms of visual category perception in human brain.

**Significance statement:** Previous works have shown where and when perceptual category information can be decoded from the human brain, our study adds to that line of research by allowing to identify spectrotemporal patterns that contribute to category decoding without the need to formulate a priori hypothesis on which spectral components and at which times are worth investigating. Application of this method to an extensive dataset of human intracerebral recordings delineates the locations that are predictive of several perceptual categories from the locations that are have narrow specialization, identifies spectral signatures characteristic of each of 8 perceptual categories and allows to observe global and category-specific patterns of neural activity pertinent to functional perceptual categorization.

## Introduction

Our capacity to categorize sensory information allows us to quickly process and recognize complex elements in our environment. Early studies revealed strong relations between brain activity within certain localized networks and neural representations of certain stimulus categories, as for example faces, bodies, houses, cars, objects and words (Kanwisher et al., 1997; Epstein et al., 1999; Peelen et al., 2009; Malach et al., 1995; Haxby et al., 2001; Ishai et al., 1999; Cohen et al., 2000). These early assessments also revealed brain networks capability to rapidly extract categorical information from short exposure to natural scenes (Potter and Faulconer, 1975; Thorpe et al., 1996; Li et al., 2002) based on models of parallel processing across neural networks (Rousselet et al., 2002; Peelen et al., 2009). In both animal and human studies, visual cortices and particularly inferior temporal cortex (ITC) appears as a key region to integrate information at the object-level (Grill-Spector and Weiner, 2014). In humans, a great deal of observations of cortical response selectivity have been achieved using fMRI, but measuring direct neuronal activity (Quiroga et al., 2005; Kreiman et al., 2000) also revealed similar patterns. To further understand how stimulus features and perceptual experience is processed in neural networks, brain activity, especially in sensory cortices, has been decoded using a variety of methods and signals (Haynes and Rees, 2006; Kriegeskorte et al., 2006; Kamitani and Tong, 2006). This decoding often relies on machine learning to avoid a priori selection of partial aspects of the data by the human observer, and unless additional analysis is performed on the model itself it does not emphasize the mechanisms of neuronal communication within and between neural networks involved in this processing.

A pervasive feature of electrophysiological neural activity are its spectral fingerprints. Neural oscillations have been proposed to reflect functional communication processes between neural networks (Fries, 2009; Buzsaki, 2006; Siegel et al., 2012). Certain frequency bands are selectively associated with the operating of different cognitive processes in the human and animal brain, (Vidal et al., 2006; Wyart and Tallon-Baudry, 2008; Jensen and Mazaheri, 2010; VanRullen, 2016; Engel and Fries, 2010; Dalal et al., 2011) and lately, direct recordings from the human cortex have revealed the remarkable representation selectivity of high gamma-band activity (50 – 150 Hz) (Lachaux et al., 2012; Parvizi and Kastner, 2018; Fox et al., 2018). Human intracranial recordings have previously shown evidence of functional processing of neural networks related to perceptual category representation (McCarthy et al., 1997) and lately the prominence of broadband high-gamma activity in selective category responses in visual areas (Vidal et al., 2010; Davidesco et al., 2013; Hamamé et al., 2014; Privman et al., 2007; Fisch et al., 2009). Yet, very little is known about the specific relation between the different components of the full power-spectrum, including high-gamma activity, and their level of selectivity in processing perceptual categories. Previous works have shown where and when perceptual category information can be decoded from the human brain, the approached introduced in this work adds to that line of research by allowing to identify spectrotemporal patterns that contribute to category decoding without the need to formulate a priori hypothesis on which spectrotemporal regions of interest are worth investigating.

In this work we capitalize on an extensive dataset of deep intracranial electrical recordings on 100 human subjects to decode neural activity produced by 8 different stimulus categories. We analyzed the decoding model built by a random forest classifier to disentangle the most informative components of the time-frequency spectrum related to the simultaneous classification of 8 different perceptual categories. Focusing on feature importance allowed us to identify activity patterns that were either characteristic of a specific visual category or were shared by several categories. In addition to feature importance we analyzed the predictive power of each activity pattern and identified how informative was their spectral signature for the classification of visual categories. In particular we tested the predictive power of high broadband gamma activity in comparison to lower frequency activity as they reflect different communication mechanisms elicited by networks seemingly involved in distinct temporal windows of functional neuronal processing. Through the analysis of feature importance of a random forest classifier we show the specific neuronal spectral fingerprints from highly distributed human cortical networks elicited during efficient perceptual categorization.

## Methods

### Patients and recordings

100 patients of either gender with drug-resistant partial epilepsy and candidates for surgery were considered in this study and recruited from Neurological Hospitals in Grenoble and Lyon (France). All patients were stereotactically implanted with multi-lead EEG depth electrodes (DIXI Medical, Besançon, France). All participants provided written informed consent, and the experimental procedures were approved by local ethical committee of Grenoble hospital (CPP Sud-Est V 09-CHU-12). Recording sites were selected solely according to clinical indications, with no reference to the current experiment. All patients had normal or corrected to normal vision.

#### Electrode implantation

11 to 15 semi-rigid electrodes were implanted per patient. Each electrode had a diameter of 0.8 mm and was comprised of 10 or 15 contacts of 2 mm length, depending on the target region, 1.5 mm apart. The coordinates of each electrode contact with their stereotactic scheme were used to anatomically localize the contacts using the proportional atlas of Talairach and Tournoux (Talairach and Tournoux, 1993), after a linear scale adjustment to correct size differences between the patients brain and the Talairach model. These locations were further confirmed by overlaying a post-implantation MRI scan (showing contact sites) with a pre-implantation structural MRI with VOXIM^®^ (IVS Solutions, Chemnitz, Germany), allowing direct visualization of contact sites relative to brain anatomy.

All patients voluntarily participated in a series of short experiments to identify local functional responses at the recorded sites (Vidal et al., 2010). The results presented here were obtained from a test exploring visual recognition. All data were recorded using approximately 120 implanted depth electrode contacts per patient with a sampling rate of 512 Hz. Data were obtained in a total of 11321 recording sites.

#### Stimuli and task

The visual recognition task lasted for about 15 minutes. Patients were instructed to press a button each time a picture of a fruit appeared on screen (visual oddball paradigm). Non-target stimuli consisted of pictures of objects of eight possible categories: houses, faces, animals, scenes, tools, pseudo words, consonant strings, and scrambled images. All the included stimuli had the same average luminance. All categories were presented within an oval aperture (illustrated on Figure 1a). Stimuli were presented for a duration of 200 ms every 1000 – 1200 ms in series of 5 pictures interleaved by 3 second pause periods during which patients could freely blink. Patients reported the detection of a target through a right-hand button press and were given feedback of their performance after each report. A 2 second delay was placed after each button press before presenting the follow-up stimulus in order to avoid mixing signals related to motor action with signals from stimulus presentation. Altogether, responses to 400 natural images were measured per subject.

**Figure 1:**
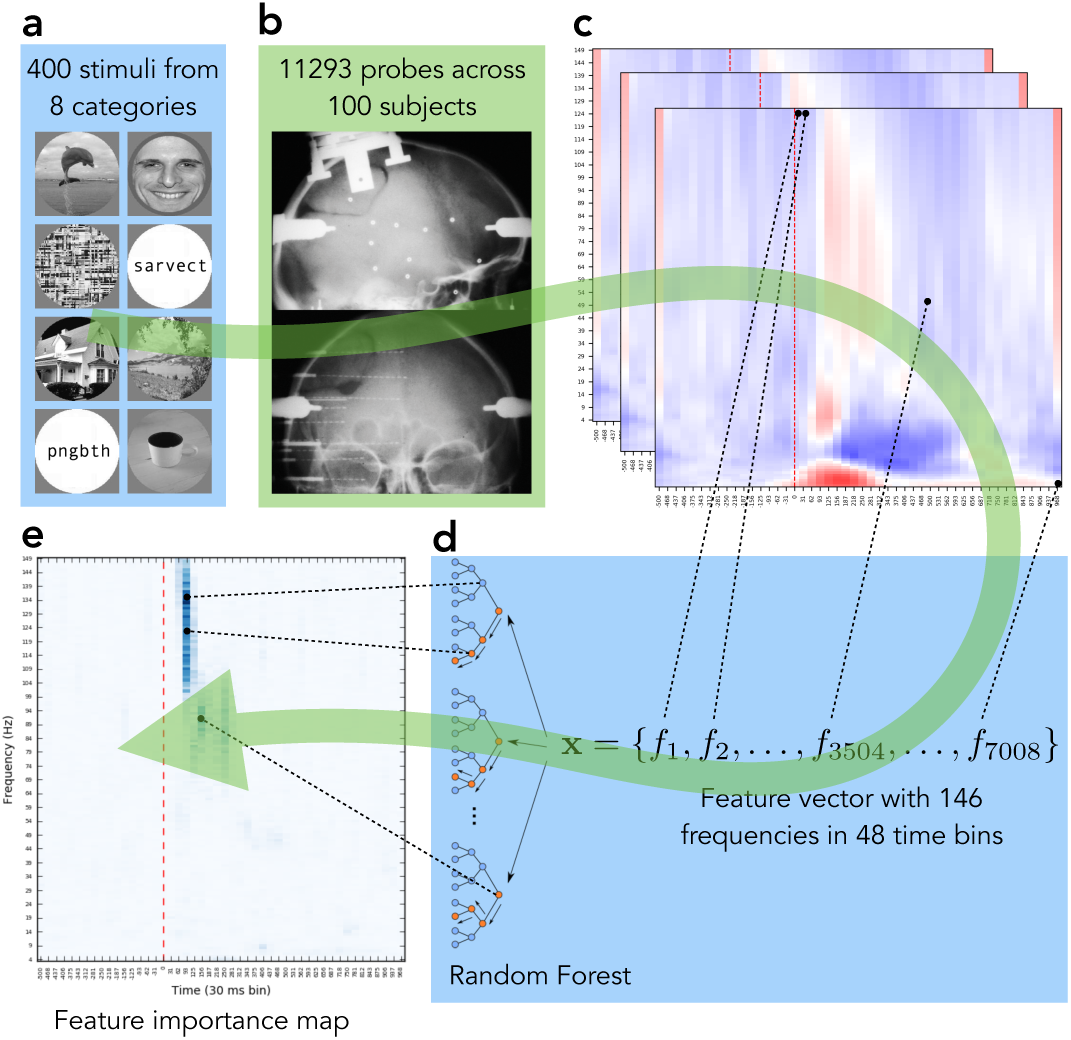
Major steps of the data processing pipeline. **a:** Image stimuli from 8 categories were presented to test subjects. **b:** Human brain responses to images were recorded with deep intracranial electrodes. **c:** LFP signals were preprocessed and transformed into time-frequency domain. **d:** Random Forest models were trained to decode image category from each electrode’s activity. **e:** Feature importances of each model were calculated to identify the region on each electrode’s activity map that was relevant to visual object recognition. Notice how the final results on panel e tell us that high gamma activity in 90 – 120 ms window and the subsequent activity in the low gamma range in 120 – 250 ms window are the only bands and time windows in that particular electrode’s activity that are relevant for the classification task, while the spectrogram on panel c also shows that there was activity in early theta, beta and low gamma bands. Our analysis revealed that not all activity was relevant (or useful) for the classification of an object and showed which parts of the activity are actually playing the role in the process.

### Processing of neural data

The analyzed dataset consisted of 4528400 local field potential (LFP) recordings – 11321 electrode responses to 400 stimuli. To remove the artifacts the signals were linearly detrended and the recordings that contained values ≥ 10*σ_images_*, where *σ_images_* is the standard deviation of responses (in the time window from −500 ms to 1000 ms) of that particular probe over all stimuli, were excluded from data. All electrodes were re-referenced to a bipolar reference. The signal was segmented in the range from −500 ms to 1000 ms, where 0 marks the moment when the stimulus was shown. The −500 to −100 ms time window served as a baseline.

To quantify the power modulation of the signals across time and frequency we used standard time-frequency (TF) wavelet decomposition (Daubechies, 1990). The signal *s*(*t*) was convoluted with a complex Morlet wavelet *w*(*t*, *f_0_*), which has Gaussian shape in time (*σ_t_*) and frequency (*σ_f_*) around a central frequency *f_0_* and defined by *σ_f_* = 1/2π*σ_t_* and a normalization factor. To achieve good time and frequency resolution over all frequencies we slowly increased the number of wavelet cycles with frequency (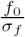 was set to 6 for high and low gamma, 5 for beta, 4 for alpha and 3 for theta frequency ranges). This method allowed to obtain better frequency resolution than applying a constant cycle length (Delorme and Makeig, 2004). The square norm of the convolution results in a time-varying representation of spectral power, given by: *P*(*t*, *f_0_*) = |*w*(*t*, *f*_0_)*s*(*t*)|^2^. Baseline normalization was performed by dividing the signal in each frequency by the average power of that frequency in the baseline window −500 to −100 ms. Each LFP recording was transformed from 768 data points (1.5 seconds of voltage readings at 512 Hz sampling rate) into a matrix of size 146 × 48 where each row represents a 1 Hz frequency band from 4 Hz to 150 Hz and columns represent 31.25 ms time bins. Value in each cell of that matrix is the power of that specific frequency averaged over 16 time points.

Further analysis was done only on the electrodes that were responsive to the visual task. In each frequency band we compared each electrode’s average post-stimulus band power to the average baseline power with a Wilcoxon signed-rank test for matched-pairs. Only the probes that showed a post-stimulus response that is statistically significantly (p-value ≤ 0.005) different from the baseline response in at least two frequency bands were preserved for future analysis. All p-values from this test were corrected for multiple comparisons across all electrodes with a false discovery rate (FDR) procedure (Genovese et al., 2002).

To anatomically localize the source of each signal in subject’s brain each electrode’s MNI coordinates were mapped to a corresponding Brodmann brain area (Brodmann, 1909) using Brodmann area atlas from MRICron (Rorden, 2007) software.

To confirm that probe’s predictiveness of a certain category implies that the probe belongs to the network selective of that category we ran a set of experiments on three well-known functional areas: Fusiform Face Area (FFA) (Kanwisher et al., 1997), Visual Word Form Area (VWFA) (Cohen et al., 2000) and Parahippocampal Place Area (PPA). Following Montreal Neurological Institute (MNI) coordinates of FFA reported in (Harris et al., 2012) and (Axelrod and Yovel, 2015) we defined FFA bounding box as *x* ∈ [−44, −38], *y* ∈ [−61, −50], *z* ∈ [−24, −15] in the left hemisphere and *x* ∈ [36, 43], *y* ∈ [−55, −49], *z* ∈ [−25, −13] in the right hemisphere. Based on the Table 1 from (Price and Devlin, 2003) we defined VWFA area as MNI bounding box *x* ∈ [−50, −38], *y* ∈ [−61, −50], *z* ∈ [−30, −16] in the left hemisphere. From MNI coordinates reported in (Bastin et al., 2013) and (Park and Chun, 2009; Hamamé et al., 2013) we defined PPA bounding box to be *x* ∈ [−31, −22], *y* ∈ [−55, −49], *z* ∈ [−12, −6] in the left hemisphere and *x* ∈ [24, 32], *y* ∈ [−54, −45], *z* ∈ [−12, −6] in the right hemisphere.

### Random Forest as a decoding model

A Random Forest (Breiman, 2001) is a collection of decision trees, where each tree gets to operate on a subset of features. Each tree is assigned a random set of features and it has to find the decision boundaries on those features that lead to best classification performance. At each branching point the algorithm must decide using which feature will be most efficient in terms of reducing the entropy of class assignations to the data points under current branch of the decision tree. To achieve that, the feature that is most useful will be selected first and will be responsible for largest information gain. For example, if the activity of a probe at 52 Hz at 340 ms is high when a subject is presented with a face and low for all other categories, decision tree will use that fact and rely on the “52 Hz at 340 ms” feature, thus assigning it some importance. How high the importance of a feature is will depend on how well does this feature distinguish faces from all other categories. As Random Forest is a collection of trees and the same feature will end up being included into several different trees, being important in many trees contributes to the overall importance of a feature (for the exact computation see the section on feature importance below).

We treated each electrode’s responses as a separate dataset consisting of 400 data points (one per stimulus image), and 7008 features – time-frequency transformation of LFP response into 146 frequencies and 48 time bins. For each electrode we trained a Random Forest with 3000 trees and used 5-fold cross-validation to measure the predictive power of the neural activity recorded by each of the electrodes. Per-class F1 score, a harmonic mean of precision and recall of a statistical model, provides us with a metric of success of the classification. The parameters were selected by performing informal parameter search. Random Forest was the algorithm of choice for our analysis due to interpretability of the resulting models, that allowed us to track the process that led each particular model to a decoding decision and due to its previous application to spectrotemporal features (Westner et al., 2018). We used `scikit-learn` (Pedregosa et al., 2011) implementation of the above-mentioned methods.

As the first step of the decoding analysis we estimated which of 11321 electrodes have predictive power. For that we split each electrode’s 400-sample dataset into 320 samples for training and 80 for prediction estimation. Repeating this procedure 5 times provided us with 400 predictions that we could compare to the true categories. By running a permutation test 100000 times on electrodes with randomly permuted class labels we estimated that 99.999th percentile (equivalent to significance threshold of *p* ≤ 0.00001) of F1 score is 0.390278. In total 787 electrodes had a predictive power of F1 > 0.390278 in at least one of the categories. For each of those electrodes a Random Forest model was retrained once more on whole data (400 samples instead of 320) and that model was used for calculating feature importances and, ultimately, for understanding which parts of the recorded activity were relevant for visual object recognition in human brain.

### Feature importance for the analysis of task-relevant neural activity

During the process of constructing the decision trees Random Forest relies on some features more than on the others. We chose *Gini impurity* (Breiman, 2017) as a measure of which feature should be used to make the branching decisions in the nodes of a tree. This score, along with the number of times each particular feature was used across trees, informed us on the relative importance of each particular feature with respect to other features. Gini impurity *G* is calculated as

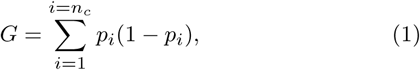

where *n_c_* is the number of categories and *p_i_* is the proportion of class *i* in a node. To pick a feature for a parent node Gini impurity of both children nodes of the parent node are calculated. The feature that decreases impurity the most is selected to be the branching factor of that parent node. The reduction in impurity is calculated as

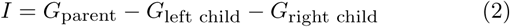

and is called *node importance. Feature importance* of feature *f* is estimated by calculating the sum of Gini impurity reductions over all samples in the dataset that were achieved with the use of a particular feature *f* and normalizing it by the total number of samples. Figure 1e is a visual representation of relative feature importance, color intensity shows the importance of each of 7008 (146 frequencies ×48 time bins) spectrotemporal features from one probe. In total our analysis has produced 787 × 8 such images − one for each probe-class pair.

The importance map computed as depicted on Figure 1 is a global map for all 8 categories, such that the regions that are highlighted on the map are important for distinguishing between all 8 categories. There is, however, a way to look at category-specific importances as well. The final set of nodes of a decision tree, called *leaves,* are the end-points of the classification process and each leaf is associated with a certain category, for example, if we take one TF activity map and start traversing a decision tree following the rules set by the nodes of the tree, we will end up in a certain leaf. And that leaf will be associated with a certain category, for example, with faces. The fact that we followed the rules and ended up in that leaf indicates that the TF map we used as the input to the tree probably comes from a trial where a “face” stimulus was shown to the subject. In order to get category-specific importance map we took all the leaves associated with a category, traverse the tree backwards and track all the features that were used on the path from the leaf to the root of the tree. This way we got a list of features that were used to classify a response to a certain category as such. Random Forest feature importance allowed us to identify which sub-regions of neural activity are relevant for decoding. It also showed that only small portion of activity is actually crucial for discrimination between the categories.

To compare importance maps between each other we fit a normal distribution on the difference between two maps and considered statistically significant the differences that are bigger than *μ* + 4*σ.* One spectrotemporal importance map consists of 7008 values. To filter out false positives we stipulated that only 1 false positive out of 7008 pixels can be tolerated and tuned the threshold accordingly. That requirement resulted in the p-value of 0.0001427 and confidence level of 99.99%, corresponding to 3.89*σ*, which we rounded up to *σ* = 4.0.

### Hierarchical clustering to reveal types of activity patterns

To further analyze the spectrotemporal signatures elicited by different visual categories in different parts of human brain we clustered filtered activity patterns and identified the most prominent groups. The result of this analysis is shown in the second column of Figure 7. For each category there are four clusters show the most common activity patterns elicited by stimuli from that category.

To do the clustering we first took each probe’s category-specific activity separately by averaging probe’s responses to 50 images of each particular category in time-frequency domain. We then masked the activity with the category importance map (as shown on Figure 3), leaving only those features out of 146 × 48 that have importance score larger that *μ* + *σ,* where *μ* is the average importance score for that category and σ is one standard deviation of the score distribution.

Masked activity patterns were hierarchically clustered using equation 3 to calculate the distance between a pair of clusters *U* and *V* as the maximal cosine distance between all of the clusters’ member observations (complete linkage clustering):

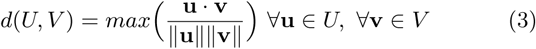

`SciPy`(Jones et al., 2001) implementation of the hierarchical clustering methods was used in this work. Resulting clustering assignments were visually inspected and corrected.

## Results

### Feature importance allows to dissociate the neural signals that are predictive of perceptual categorization from the rest of the stimulus-induced neural responses

To identify spectrotemporal features that are characteristic of perceptual categorization of a particular category we relied on time-frequency (TF) maps of the neural responses of intracranially implanted electrodes. Out of the total set of 11321 probes 11094 (98%) were responsive (see the Methods section on processing of neural data for details) to the stimuli from at least one of the categories. On one hand this provides us with abundance of data, on the other raises the question whether all of that activity was relevant to the processes that encode and process visual input.

Training a decoding model (see the Methods section on Random Forest as decoding model) for each of the probes allowed us to dissociate the *predictive probes* that exhibited activity that was useful for decoding from the rest of the *responsive probes* that did not carry such activity.

Green markers on Figure 2a show the set of probes that are responsive to the `house` category, while the blue markers are the probes that are predictive of that category (4.8%, 535 probes). Decoding models built on the neural responses of the predictive probes were successful at classifying at least one perceptual category, focusing on them in our further analysis allowed to work only with the locations that carry information relevant to the task of perceptual categorization.

**Figure 2:**
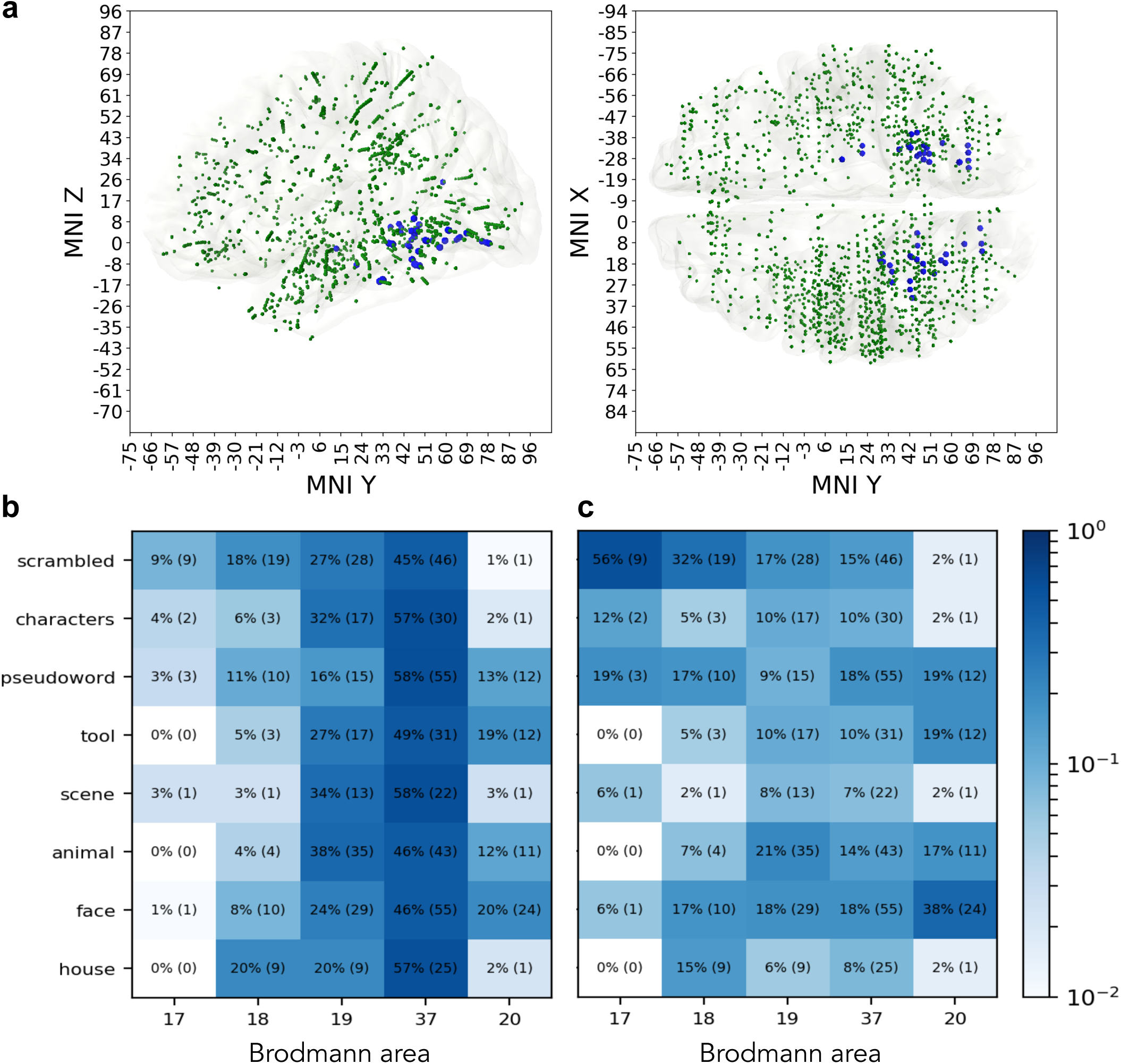
Distribution of predictive probes. **a.** Green markers indicate the probes that were responsive to stimuli from the `house` category. Blue markers are the predictive probes that carry information that is relevant to decoding the neural response as reaction to `house` stimulus. **b.** Distribution of predictive probes over areas within each category. **c.** Distribution of predictive probes over a category with each area.

Predictive probes had heterogeneous distribution in the brain, yet remained mostly concentrated in visual cortices and inferior temporal regions (76%), from BA17 to BA20, including early visual areas (BA 18, 19), fusiform gyrus (BA 37) and inferior temporal cortex (BA 20). A majority of the predictive probes were in fusiform cortex (average of 52% over all categories, Figure 2b), followed by BA 19 (27%), across all category networks.

Within the primary visual cortex, BA 17 and 18, the `scrambled` (control condition) was the stimulus that elicited most predictive probes amongst all stimulus categories (Figure 2c), followed by `pseudowords` and `characters`. Probes predictive of `faces` were mostly concentrated in BA19, BA37 and BA20. The low number of predictive probes in area 17 is explained by the fact that less than 1% of the implantation sites in the original dataset were located in primary visual cortex.

Previous studies have shown that perceptual category-selective networks are located in occipito-temporal cortex (Grill-Spector and Weiner, 2014; Ishai et al., 1999; Malach et al., 1995). To test whether predictive power computed by the Random Forest model trained to decode activity of probes is coherent with known functional processing by cortical networks we evaluated the selectivity of the predictive power in three known functional networks: Fusiform Face Area (FFA) (Kanwisher et al., 1997), Visual Word Form Area (VWFA) (Cohen et al., 2000) and Parahippocampal Place Area (PPA) (Epstein and Kanwisher, 1998). We checked whether the probes located in each of these areas and the Random Forest model trained on these probe’s activity to discriminate between 8 categories produces the highest predictive power for the category for which this area is known to be selective. Probes in FFA are expected to be good at discriminating `faces`, probes in VWFA should be predictive of `characters` and `pseudowords` categories and probes in PPA should be responsive to `scenes` and `houses`.

There were 12 probes in the FFA that were predictive of a category: 5 were predictive of `faces`, 4 of `animals` (which mostly have faces on the image), 2 of `pseudowords` and 1 of `scrambled` images. Most probes that were in FFA and were predictive, carried information of the categories containing facial features.

There were 8 probes in the VWFA that were predictive of a category: 5 were predictive of `pseudowords`, 2 of `characters` and 1 of `faces`. This points to the fact that the predictive probes in VWFA are predictive of the stimuli with written characters on them. These results confirm that predictive power of a Random Forest model trained on probes activity in VWFA reflects the functional role known to be carried by this area.

For probes in the PPA results were less selective. There were 23 probes inside that area that were predictive of a category: 5 were predictive of `houses`, 4 of `scenes`, 5 of `characters`, 5 of `scrambled` images, 2 of `tools` and 2 of `pseudowords`. Here the case for functional role of the area is less straightforward as the probes from PPA predicted not only `houses` and `scenes`, but also other categories. However, `houses` and `scenes` were among the categories that the probes from PPA were able to identify successfully in highest proportion as compared to the other categories.

We find that these confirmatory findings give credibility to the fact that the probes that are predictive of a certain category according to our method are indeed involved in the processing of the stimuli that belong to that category.

Training per-probe decoding models not only allowed us to identify the predictive locations, but also to apply feature importance analysis to decoding models trained on local activity. Computing the feature importance across the time-frequency map (4 – 150 Hz and −500 to 1000 ms) allowed us to see which part of neural activity is crucial for in the decoding. Overlaying the importance over time-frequency map showed at which frequencies and at what times the activity that was important for the algorithm has occurred. This can be done both on aggregated level, where the importance map is averaged over probes, and on individual probe level. Figure 3 illustrates the application of probe importance map to filter irrelevant activity and obtain spectrotemporal signature of a particular category on a particular probe. Now we can use the feature importance map as a mask and dive into the analysis of the activity itself, but focus only on the relevant parts of it. We believe that, when applicable, this methodology helps to filter out irrelevant activity and allows to focus on the activity that is important to the scientific question under investigation.

**Figure 3:**
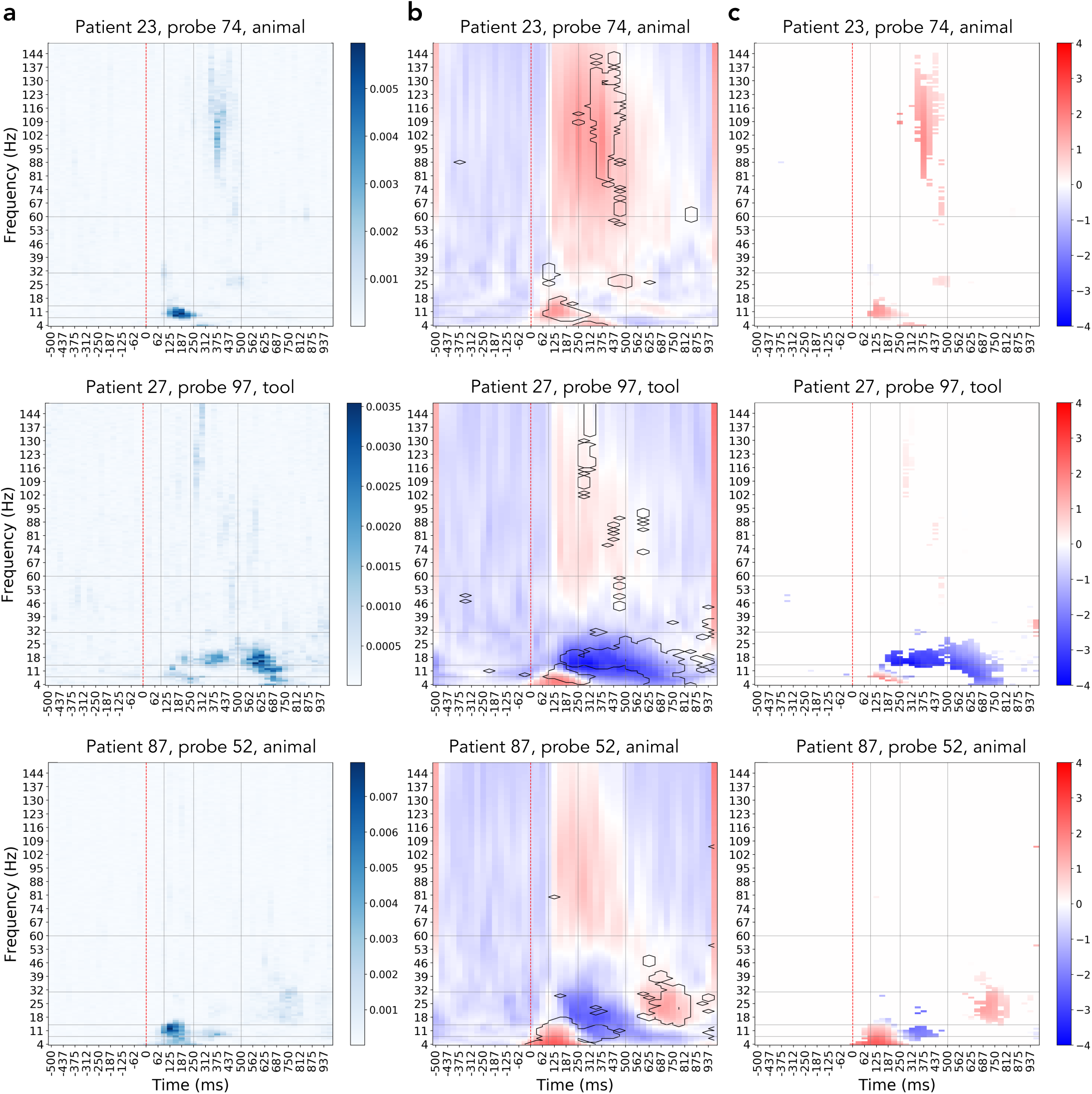
Using the importance map to filter out irrelevant activity. The three rows show three different examples of how filtering of the activity by importance is beneficial: in patient 23, probe 74 we see that only later portion of the broadband gamma activity increase was useful for identifying this activity as a response to the `animal` stimulus; patient 27, probe 97 shows that although there is an increase in broadband activity, the actually useful information contained in decrease in the lower frequency bands; patient 87, probe 52 demonstrates that for decoding this particular probe’s activity one must focus on the activity in lower frequencies at specific time and, despite prominent presence, ignore the increase in broadband gamma. **a.** Probe’s importance map, color codes the relative importance of each spectrotemporal feature within the map. **b.** Full spectrotemporal activity of the probe, features with importances one standard deviation higher than the average (in contour) mark the regions of activity that were useful for the decoding model. **c.** Activity of the probes filtered by the importance mask, only the relevant activity is preserved.

We took an average over importance maps of all individual probes within each category to obtain the global picture of where the category-specific activity lies in time and frequency space. Figure 4 summarizes such analysis and allows to see which spectrotemporal signatures are unique to perceptual categorization of specific categories and which ones are shared across several ones. From these importance maps we notice that certain TF components are distinctly present per category, as for example high importance of the transient theta activity (theta burst) in all categories, or the almost absence of broadband gamma in the control scrambled condition. In the following sections we expand our analysis to the comparison of the feature maps and analyzing the activity under the regions that we have identified as important.

**Figure 4:**
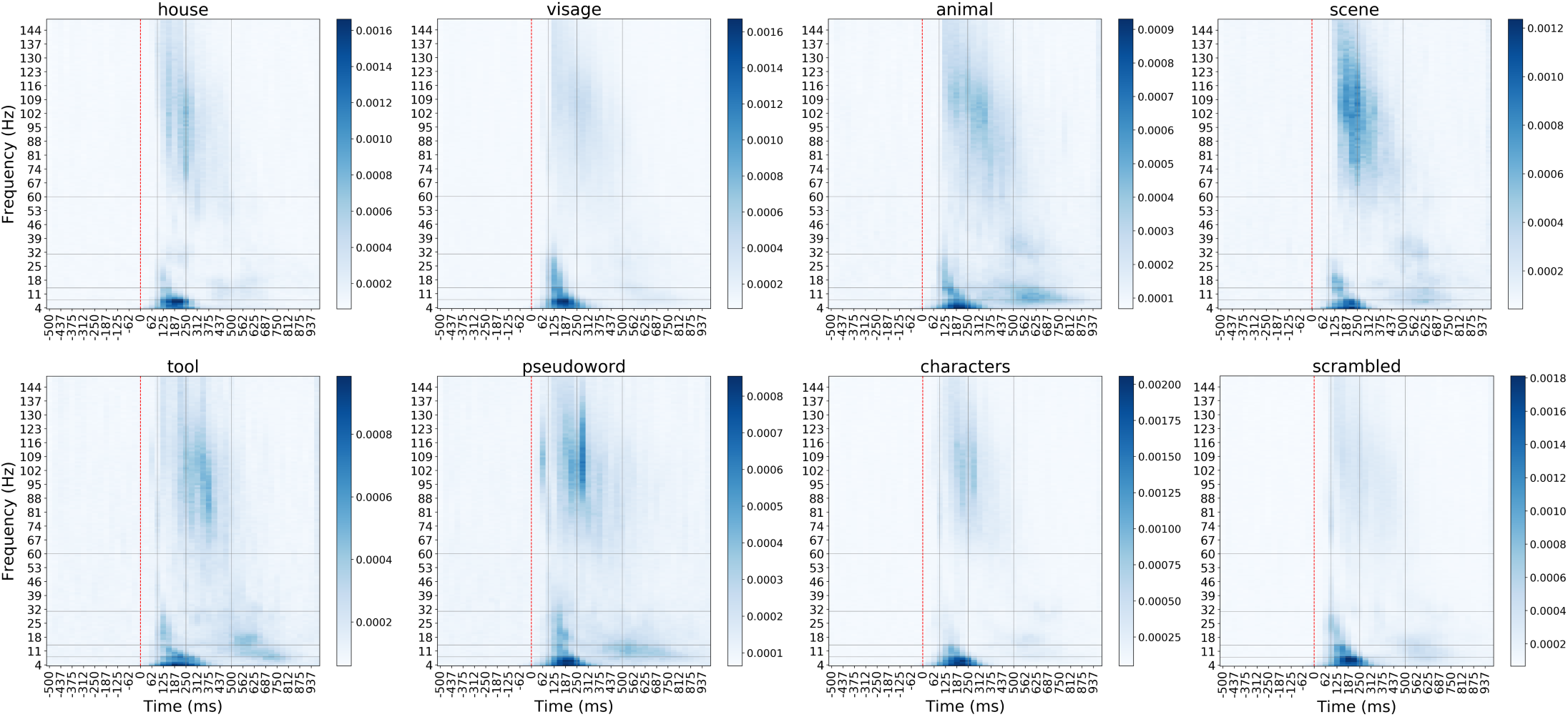
Average importance map of each of eight categories over probes predictive of that category. The color shows the relative importance of each spectrotemporal feature, indicating how informative that particular feature was for the task of decoding.

### Polypredictive and monopredictive probes

Spectral decomposed neural activity revealed two types of probes regarding the number of categories that could be decoded. From some probes it was possible to decode more than one category with above-chance classification accuracy (Figure 5b). We considered a probe to be predictive of a category if cross-validation F1 score for that category was higher than 0.39 (see the Methods section for details on threshold selection), which is a stricter condition than above-chance criterion (corresponding to F1 > 0.125). There is a subset of networks in the brain that processes information relevant to more than a single visual category, we refer to the locations that constitute those networks as *polypredictive.* Other locations, which we call *monopredictive,* are useful in predicting only one out of 8 different types of stimuli revealing a high degree of specialization. Figure 5a shows that polypredictive probes reside mainly in posterior occipital and posterior temporal cortex networks, while monopredictive probes extend, in addition to occupying similar posterior occipital and temporal locations, to frontal cortex and anterior temporal cortex. Both mono- and polypredictive probes are also observed in parietal cortex. Monopredictive probes that extend beyond ventral stream and temporal cortex pertain to the following perceptual categories: `faces` (orbitofrontal cortex), `animals` and `pseudowords` (dorsofrontal cortex, inferior frontolateral cortex, premotor cortex), and, to a smaller extent, `scrambled` images (prefrontal cortex).

**Figure 5:**
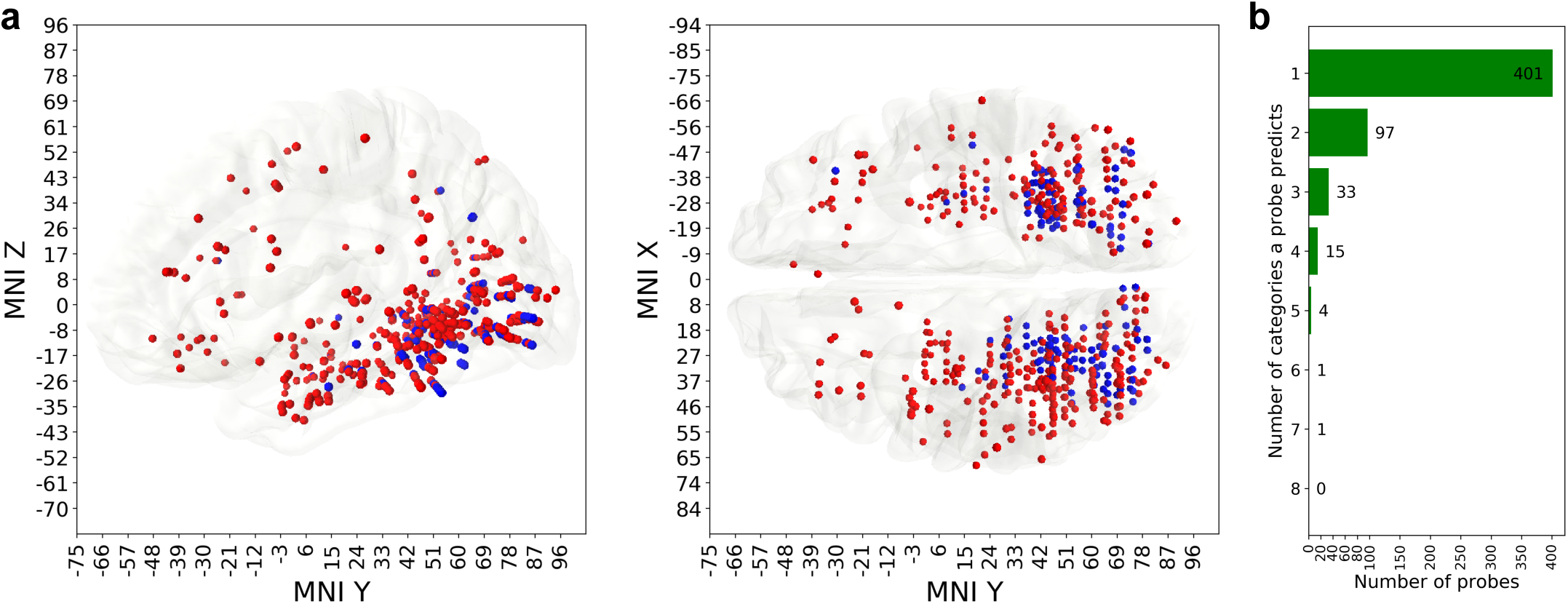
Anatomical distribution of mono- and polypredictive locations. **a:** Red markers are the locations of monopredictive probes, blue markers are the locations of polypredictive ones. Polypredictive probes (145 unique locations) are mostly confined to visual areas and temporal lobe (both parts of the ventral stream), while monopredictive (specialized, 401 unique locations) probes are, in addition to visual areas, also found in frontal and parietal cortical structures. **b:** The histogram shows how many categories are predictable by how many probes.

The unique association of specific TF feature importance components with either polypredictive and monopredictive probes was category specific, as shown in figures 6a to 6h. For `face` stimuli, most of the feature importance in the early broadband gamma response and a part of the early theta response was significantly stronger in polypredictive probes as compared to monopredictive probes, yet present in both and predictive of the category as compared to all other categories (Figure 6b). For `face` stimuli no specific part of the TF feature importance was stronger in monopredictive probes as compared to polypredictive probes. `Animals` and `tools` categories showed a prevalence of feature importance of monopredictive activity patterns in late broadband gamma range (> 300 ms) and in the alpha/beta range, with very little dominant feature importance TF components in polypredictive probes. `Scenes` and houses also show stronger feature importance in late alpha and beta band responses in monopredictive probes. Interestingly, for `characters` (Figure 6g), feature importance in the early broadband gamma range was dominant for polypredictive probes, while for `pseudowords` (Figure 6f) the late broadband gamma revealed to be dominant for monopredictive probes. `Pseudowords` also elicited a significantly stronger TF feature importance in monopredictive probes in the low-frequency range, similar to `animal` and `tool` stimulus categories. Finally, an interesting observation was that `animals` and `faces` share most of their polypredictive probes (more than 50%) indicating a large overlap of categorization networks of these two categories.

**Figure 6:**
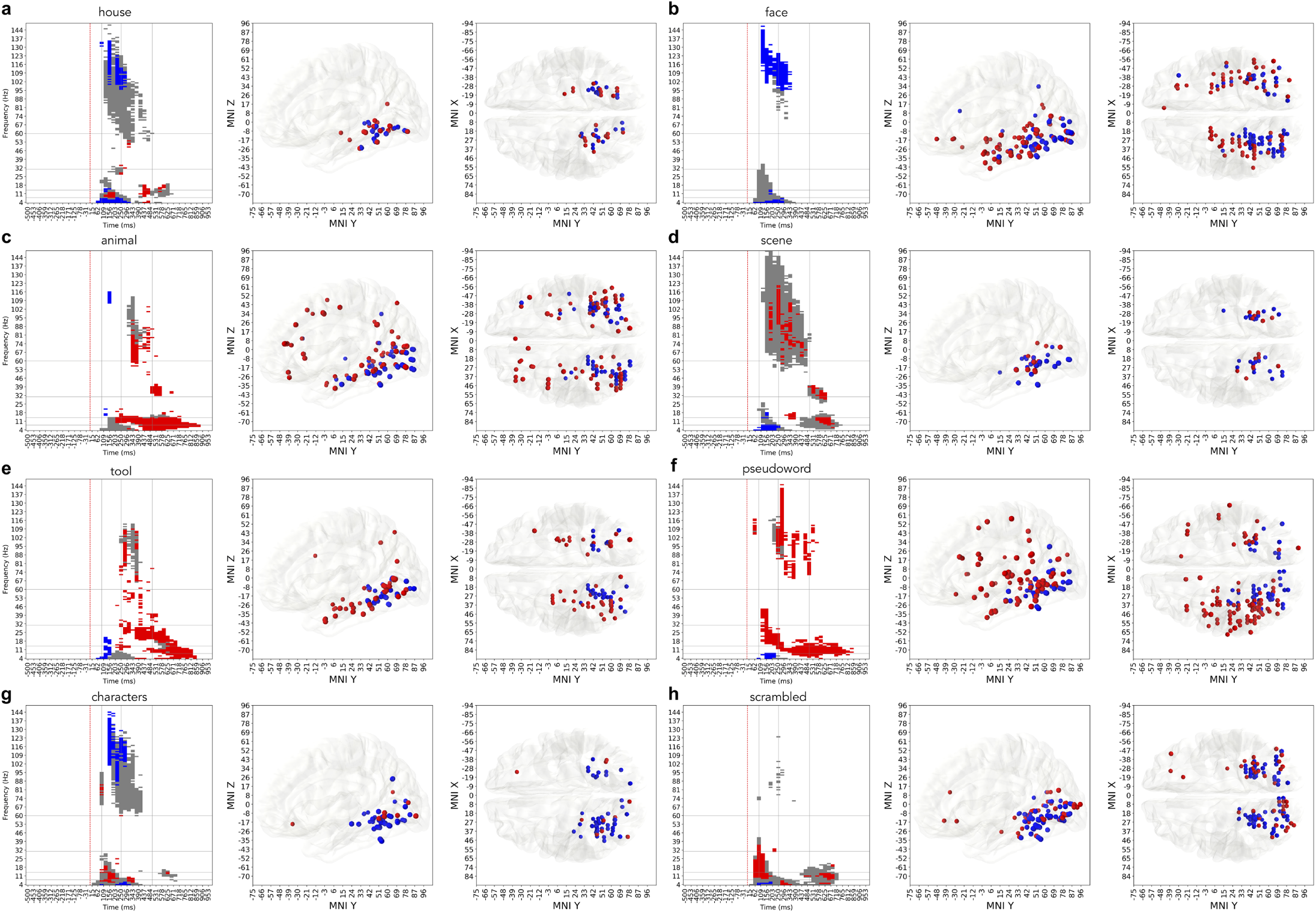
Difference between time-frequency feature importances and locations of monopredictive (red) and polypredictive (blue) probes for each category. **a:** `house`, **b:** `face`, **c:** `animal`, **d:** `scene`, **e:** `tool`, **f:** `pseudoword`, **g:** `characters`, **f:** `scrambled`.

### Diversity of time-frequency patterns plays a role in functional perceptual categorization not only across but also within each perceptual category

To quantify which TF components group together within each category network across the probes predictive of that category we clustered the probes according to their activity masked by feature importance. Left column of figure 7a shows an averaged feature importance map for a given category. Next, we look into the regions of time-frequency map that are indicated as important by the feature importance map, extract baseline normalized activity in those regions and cluster that activity using hierarchical complete linkage clustering with cosine distance (see the Methods section on hierarchical clustering for details). The second column of figure 7 shows the activity of four most populated clusters for each category. Each cluster represents the activity pattern exhibited by the probes in that cluster. Only the probes whose activity had predictive power are included in this analysis. As the final step we identified the anatomical locations of the probes from each cluster to see whether difference in the activity patterns could be attributed to the functional regions of the brain. The visualization of this step in the last two columns of figure 7.

**Figure 7:**
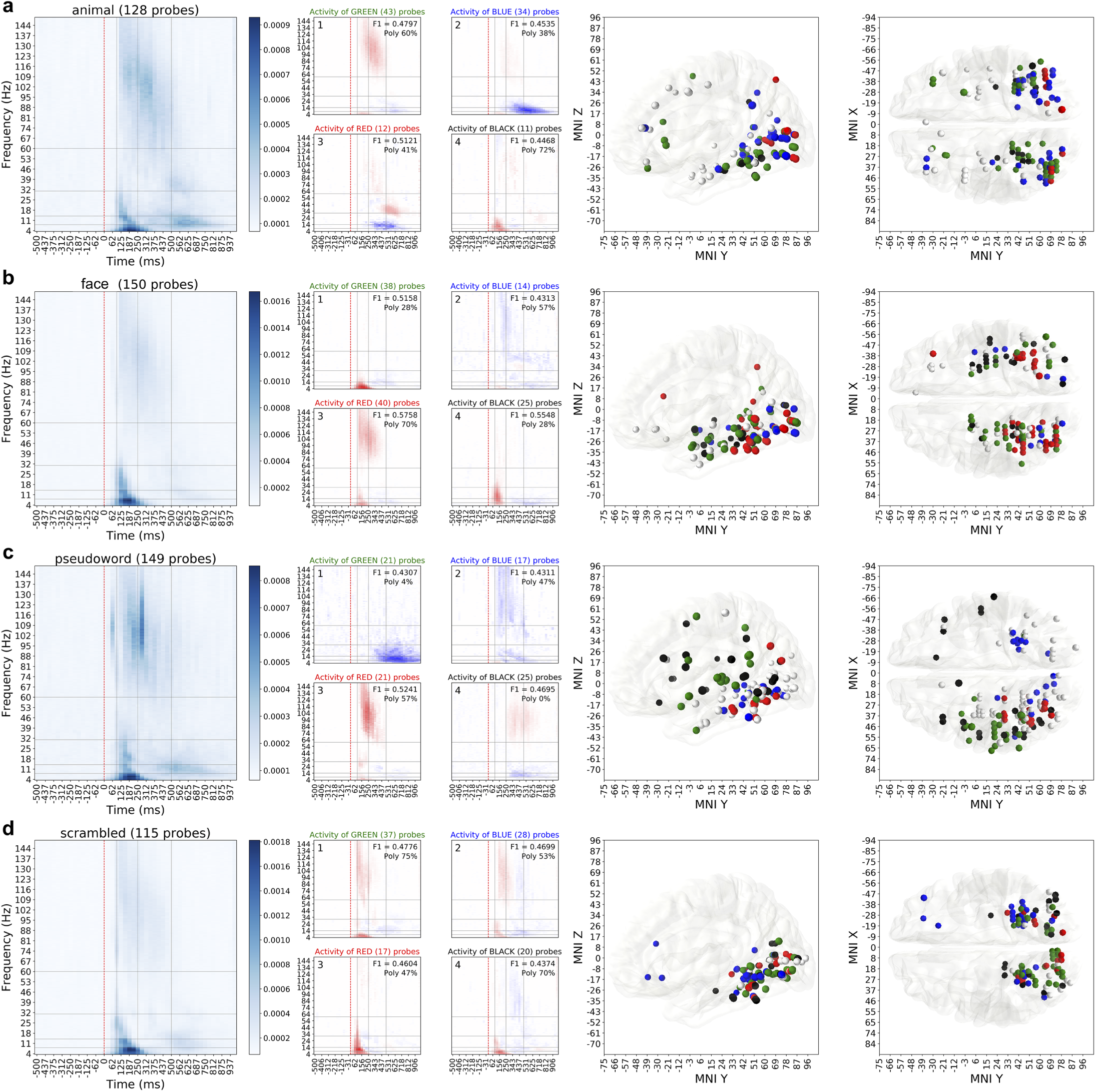
Detailed analysis of spectral activity of **(a)** `animals`, **(b)** `faces`, **(c)** `pseudowords` and **(d)** `scrambled` images. Leftmost column contains the importance maps extracted from Random Forest models and shows where in time and frequency the important activity is. Second column visualizes the four largest (by the number of recording sites) clusters of activity patterns inside those spectrotemporal regions that are deemed important. The numbers in the top right corner of each cluster’s activity pattern show the average predictive power (F1 score) of the probes in that cluster and proportion of polypredictive locations that exhibited this particular pattern of activity. Note how every cluster has a designated color: green, blue, red or black. This color of the cluster matches the color of MNI location markers in the last two columns, that show sagittal and dorsal views of the brain. White markers show the probes that have predictive power, but their activity pattern does not belong to any of the four major clusters.

This analysis allowed us make a number of *global,* pertaining to the categorization of visual stimuli over all categories, as well as number of *category-specific* observations.

The first global observation was that it is not only broadband gamma activity that is useful for the random forest model performance, but low-frequency activity also contributed significantly sometimes overshadowing the activity of higher frequency bands altogether. Most clusters were composed of a combination of low and high-frequency components (Figure 7, second column) and were mostly located in occipito-temporal cortices, though some electrodes in parietal and frontal cortex also appeared to contribute with significant responses in the classification (Figure 7, two right columns), especially for such stimulus categories as `animal` and `pseudoword`.

The second observation spanning across all categories was that the classifier used not only the increases in power to perform the classification, but also relied on power decreases in different brain networks. The most prominent examples are the clusters `faces-2` (Figure 7b), `animals-2` (Figure 7a), `tools-2`, `pseudowords-1`, `pseudowords-2` (Figure 7c), `characters-1` and `characters-2` (Figure 7d). For example, to decode `face` or `pseudowords` from the activity of the blue cluster network, the RF classifier used broadband gamma power decreases located in posterior inferior temporal cortex and inferior occipital gyrus. None of the probes for which the decrease in activity was identified as important for decoding were located in classically defined Default Mode Network (Buckner et al., 2008; Raichle, 2015).

Across all categories, the earliest component that often appeared in clusters was the brief power increase in the low-frequency interval (2-25 Hz), which for one group of probes can be associated to an almost instantaneous broadband gamma power increase (7b, cluster 3), but remains the only source of important activity for another group of probes (7b, cluster 1).

Studying the anatomical locations of the probes belonging to different clusters of activity revealed interesting observations. Figure 7c, `pseudowords`, clusters 1 and 3 show a clear example how clustering by activity patterns leads to assigning the probes into functionally different anatomical areas. The gamma-band increase signature captured by cluster 3 occurs only in the left hemisphere (red markers on Figure 7c), the late theta-alpha power decrease captured by cluster 1 also occurs only in the left hemisphere (green markers) and is spatially clearly distinct from probes in cluster 3. Because it is known that `pseudoword` stimuli elicit top-down language-related (phonological) analysis, which elicits highly left-lateralized networks identifiable in iEEG recordings (Juphard et al., 2011; Mainy et al., 2008), we know that this observation reflects a functional brain process. This dissociation in both the spectrotemporal and anatomical domains provides us with valuable information about the process of perceptual categorization and highlights the benefit of disentangling the activity into functionally and anatomically disassociated components.

We were also interested to study the relevance of the different components in the TF domain for the Random Forest classification process. Specifically, we wanted to see whether the activity in the broadband gamma range, commonly present on most clusters across categories, is in general the most valuable neural signature for category networks as compared to the low-frequency parts of the spectrum. To test whether broadband gamma was solely the most informative frequency interval we statistically compared predictive power of three intervals: broadband gamma (50 – 150 Hz), low-frequency (4 – 50 Hz) and full spectrum (4 – 150 Hz). Overall, across 7 perceptual categories out of 8 (except for `scenes`), using the full spectrum was more informative than using the broadband gamma interval or the low-frequency interval alone (Mann−Whitney U test, *p* < 0.001563 (0.05 Bonferroni-corrected to the number of clusters compared, see Figure 8). For `scrambled` images and `faces` the broadband gamma carried *less* information that was relevant for decoder than the lower frequencies.

**Figure 8:**
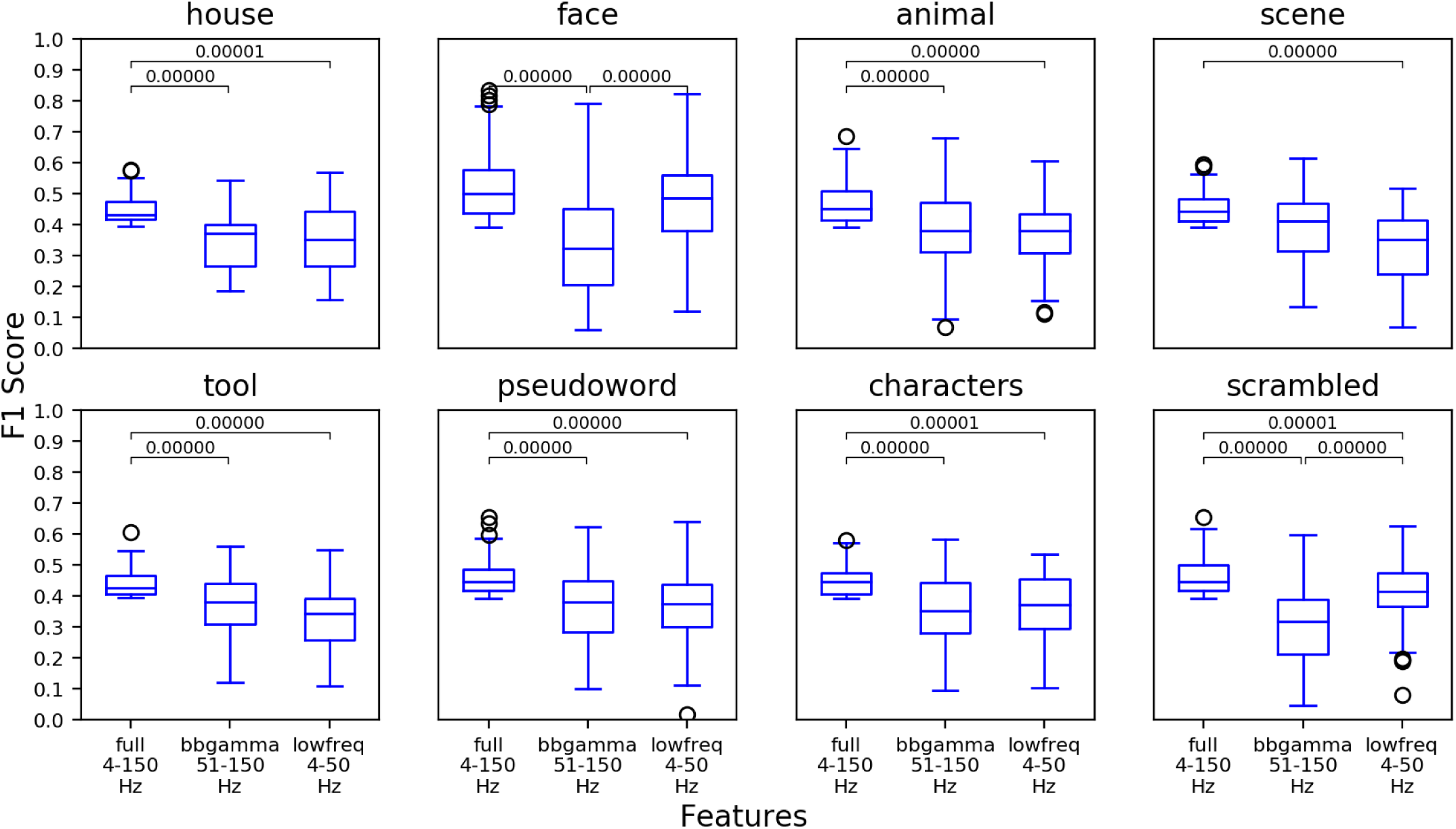
Comparison of predictive power of three different sets of features: full spectrum (4 – 150 Hz), broadband gamma alone (50 – 150 Hz) and lower frequencies alone (4 – 50 Hz) across categories. The bracket with the p-value indicates a significant difference according to Mann-Whitney U test.

## Discussion

In the present work we trained a machine learning model to decode 8 different perceptual categories from human intracerebral neural spectral activity and analyzed the resulting statistical model to expose what information does the model rely on in order to make decoding decisions. We then drew the parallels between the patterns of neural activity that the machine learning algorithm deemed important and the functional role of that activity for the task of decoding perceptual categories in human brain. This allowed us to distinguish between the spectral activity that is relevant for the task from the activity that is not. The study was conducted using a large dataset consisting of 2823250 local field potential recordings, from a cohort of 100 human patients whose combined electrode implantations spanned across all major brain cortical areas. The decoding process confronted neural responses elicited by 8 different perceptual categories (from 8 different stimulus sets), which allowed obtaining a high resolution in the dissociation between neural response patterns across categories. All classifications were operated on a broad frequency spectrum ranging from 4 to 150 Hz and allowed to distinguish degrees of selectivity of neural responses and which spectral components most strongly enable this selectivity. Previous works have shown where and when perceptual category information can be decoded from the human brain, our study adds to that line of research by allowing to identify spectrotemporal patterns that contribute to category decoding without the need to formulate a priori hypothesis on which spectral components and at which times are worth investigating.

The classifier model first allowed us to globally identify two types of neural responses: those that were predictive of a certain category and those that did not predict any category despite eliciting strong amplitude modulation across multiple frequency bands. Surprisingly, when comparing the level of predictability of probe responses we found that only 4.8% of the responsive probes were predictive of a category. This very low percentage highlights an important fact regarding the level of “selectivity” of a neural responses. When comparing neural responses, elicited by a specific stimulus, to a single or few other stimulation conditions it might seem to be distinct and thus selective, yet, when increasing the number of comparisons to a broader variety of response conditions, the degree of its selectivity might narrow down as its similarity to other responses increases. Stimulus-induced neural signal selectivity is thus a graded quality that can be assessed through multiple comparisons with a broad variety of stimulation conditions. This result also implies that although any stimulus can elicit a local neural response throughout the cerebral cortex, in the light of our results, there is a high probability of it being non-predictive of any of the categories or being polypredictive of several categories at once.

In line with a vast literature on the localization of category related networks (Kanwisher et al., 1997; Epstein et al., 1999; Malach et al., 1995; Haxby et al., 2001; Ishai et al., 1999; Grill-Spector and Weiner, 2014; Cohen et al., 2000; Peelen et al., 2009) predictive probes concentrated mostly in the inferior temporal cortex, namely the fusiform gyrus (BA 37), yet surprisingly for some categories, probes in primary visual cortex were also predictive of these categories. This effect is probably related to the specifics of the physical content of certain images that uniquely characterize certain categories amongst all others, as for example the content in high-contrast edge information in `scrambled` and written text stimuli.

Predictive probes were subsequently classified according to their level of selectivity towards a single or multiple visual categories. Polypredictive probes (36%) clustered in visual cortices and inferior temporal cortex and were associated with early spectral components (< 300 ms) such as broadband gamma power increases and a transient theta burst shortly after stimulus presentation. Monopredictive probes (64%) were abundant in these same regions, but extending uniquely in frontal, parietal, superior temporal and anterior limbic cortex. Their activity was strongly associated with the later (> 300 ms) time and with power suppression of spectral importance features, versus baseline, in the theta (4 – 7 Hz), alpha (8 – 15 Hz) and beta bands (16 – 40 Hz). In a subgroup of probes the associated power suppression of the feature importances extended into the broad gamma band (50 – 150 Hz).

Importantly, the capacity to ascribe category selectivity to predictive probes (mono vs polypredictive probes) arises from the fact that the decoding model was trained to discriminate between all 8 categories simultaneously. The separation between mono and polypredictive probes revealed specific effects in terms of network localization and time-frequency components. The high concentration of polypredictive probes (and local networks) in early visual cortices, from primary visual cortex up to inferior temporal cortex is coherent with the idea that networks in the ventral visual stream progressively integrate more complex features into object representations, thus becoming progressively more selective, and converge within median temporal lobe to more stimulus-invariant representations (Quiroga et al., 2005). This progressive information integration by spectral features of neuronal responses across the visual hierarchy has been recently connected with the computations carried out by deep convolutional neural networks trained to solve the task of visual recognition (Kuzovkin et al., 2018).

Globally, the random forest data classification provided results that are coherent with current knowledge on 1) the implication of networks located in visual cortex and inferior temporal cortex in processing visual categories, 2) the timing of object categorization in the human brain and 3) the role of broadband gamma responses in processing category-selective information within these networks. Previous studies have shown that certain stimulus categories elicit clustered cortical responses of highly localized networks in the occipito-temporal ventral stream such as the fusiform-face-area (FFA) and the visual-word-form area (VWFA) (Kanwisher et al., 1997; Cohen et al., 2000). Yet, other studies have broadened this scope by showing that certain categories, as for example faces, rely on the involvement of a larger brain-wide distributed network (Ishai et al., 2005; Vidal et al., 2010). Our classification analysis shows that the spatial extent of this network distribution is category specific, certain stimuli eliciting larger network responses, such as for `faces`, `animals` and `pseudowords`, as compared to `scenes`, `houses` and `scrambled` images which concentrate in the fusiform cortex, the parahippocampal cortex and primary visual cortex respectively.

Our results largely agree with previous works trying to decode visual object categories over time with MEG (Carlson et al., 2013; Cichy et al., 2014) or intracranial recordings (Liu et al., 2009). All these studies converge on the result that perceptual categories can be decoded from human brain signals as early as 100 ms. Our current work goes a step beyond these previous investigations by demonstrating which spectral components underlie this fast decoding. Previous intracranial studies have also shown that broadband gamma is modulated by information about object categories (Vidal et al., 2010; Privman et al., 2007; Fisch et al., 2009). Moreover, broadband gamma has been suggested as a proxy to population spiking output activity (Manning et al., 2009; Ray and Maunsell, 2011; Lachaux et al., 2012; Ray et al., 2008). It has since then been considered as a hallmark of local population processing (Parvizi and Kastner, 2018). Our classification results however show that broadband gamma is not the sole selectivity marker of functional neural processing, and that higher decoding accuracy can be achieved by including low-frequency components of the spectrum. For certain stimulus categories, as scrambled images, the broadband gamma range is even outperformed by the predictive power of the low-frequency range.

To understand which spectral components play a specific role in stimulus categorization we analyzed the decision process that drives the decoding model and identified the combined spectrotemporal regions that are informative for the output of the random forest classification procedure. This allowed us 1) to show the category-selective association of different spectral components with activity patterns represented by high feature importance, and 2) identify a functional role of positive as well as negative power modulations (increases and decreases versus baseline) in early and late time windows of neural processing involved in visual categorization.

While early TF components (i.e. broadband gamma and theta burst) appeared to reflect a polypredictive neural process across categories, especially for `faces`, the sustained decrease in power in the alpha/beta band was extended in space and time. This process is probably dependent on the degree of difficulty for the networks in reaching a perceptual decision and which appeals to the involvement of top-down processing required to resolve perceptual ambiguity elicited by the different stimulus categories. For example, `animal` and `tool` stimuli are highly diverse in their physical image structure, as compared to `face` stimuli. This affects the efficiency of bottom-up process in extracting category information, often associated with increase in gamma activity, and probably in parallel triggers top-down processes through selective activity modulation in low-frequency channels (Bastos et al., 2015). In our data, this latter phenomenon could be mirrored by a decrease of predictive power in the low-frequency range. Studies have shown that power modulations reflect changes in network connectivity (Tewarie et al., 2018) and that top-down processes, eliciting a decrease in power in the alpha-beta band, are accompanied by an increase in distant network connectivity (Gaillard et al., 2009).

Finally, we also show that certain probes elicit decreased broadband gamma responses (versus baseline) while representing a significant feature importance for the classification model. It has been shown that neural activity in the Default Mode Network can be negatively modulated by attending sensory stimulation (Buckner et al., 2008), and intracranial studies have found that this was reflected by decreases (versus baseline) in the broad gamma range (Ossandón et al., 2011; Jerbi et al., 2010; Dastjerdi et al., 2011). Here we found no evidence of such power decreases in probes located in the DMN (Buckner et al., 2008). However, the random forest classifier singled-out broad spectral patterns of power decreases at probes located in visual regions and beyond for categories `faces`, `pseudowords` and `characters`. This is the first time, to our knowledge, that power decreases in the broadband gamma range outside the DMN have been associated with highly functional neural signal classification of perceptual categories. Their functional significance should be studied in the future as they could reflect an important phenomenon of communication regulation between networks during perceptual decision making of visual categories.

In this work we studied the information that allows local cortical populations of neurons to make predictions of the category of a visual stimulus. The number of locations per patient we were able to record from was below a hundred, expanding on this work by including more subject data in the future might allow us to make a transition from the observations of local activity and the analysis of its role to being able to detect signatures of global decision-making processes. It is possible that these signatures would be reflected in specific spectral fingerprints as many classic theories would suggest (Rodriguez et al., 1999; Varela et al., 2001; Engel et al., 2001; Siegel et al., 2012). We believe that the methodology proposed in this study can facilitate the search of those fingerprints.

Our work explored which spectral components reflect the perceptual categorization process. We observed that some spectral signatures were specific to certain categories, while other were common across multiple categories. Moreover, we found that broadband gamma band is not the only or for some categories even not the main spectral component relevant for successful categorization. Intracranial recordings allowed us to see where, when and at which frequencies category information is available for the brain, hence providing fuller picture of neural processes leading up to perceptual categorization.

## Acknowledgements

IK and RV thank the financial support from the Estonian Research Council through the personal research grants PUT438 and PUT1476. This work was supported by the Estonian Centre of Excellence in IT (EXCITE), funded by the European Regional Development Fund. JA was also supported by the European Unions Horizon 2020 Research and Innovation Programme under the Marie Skodowska-Curie grant agreement no. 799411. JPL and SR received funding from the “Programme Avenir Lyon Saint-Etienne” of Universit‘e de Lyon, within the program “Investissements d’Avenir”, ANR convention No. ANR-11-IDEX-0007; the LabEx CORTEX, ANR convention No. ANR-11-LABX-0042; and European Union Seventh Framework Programme (FP7/2007-2013) under Grant Agreement No. 604102 (Human Brain Project); icEEG-only recordings were made possible thanks to IHU CESAME, within the program “Investissements d’Avenir” (ANR-10-IBHU-0003). JPL received funding from HBP Grant Agreements No. 785907 (SGA2) and No. 7202070 (SGA1). MPB received funding from the Institut Universitaire de France.

